# A small molecule ligand for the novel pain target GPR171 produces minimal reward in mice

**DOI:** 10.1101/2022.07.05.498847

**Authors:** Max V. McDermott, Akila Ram, Matt T. Mattoon, Emmaline E. Haderlie, Megan C. Raddatz, Madi K. Thomason, Erin N. Bobeck

## Abstract

ProSAAS is one of the most abundant proteins in the brain and is processed into several smaller peptides. One of which, BigLEN, is an endogenous ligand for the G protein-coupled receptor, GPR171. Recent work in rodent models has shown that a small-molecule ligand for GPR171, MS15203, increases morphine antinociception and is effective in lessening chronic pain. While these studies provide evidence for GPR171 as a possible pain target, its abuse liability has not yet been assessed which we evaluate in the current study. We first mapped the distribution of GPR171 and ProSAAS throughout the reward circuit of the brain using immunohistochemistry and showed that GPR171 and ProSAAS are localized in the hippocampus, basolateral amygdala, nucleus accumbens, prefrontal cortex. In the major dopaminergic structure, the ventral tegmental area (VTA), GPR171 appeared to be primarily localized in dopamine neurons while ProSAAS is outside of dopamine neurons. Next, MS15203 was administered to mice with or without morphine, and VTA slices were stained for the immediate early gene c-Fos as a marker of neuronal activation. Quantification of c-Fos-positive cells revealed no statistical difference between MS15203 and saline, suggesting that MS15203 does not increase VTA activation and dopamine release. Similarly, the results of a conditioned place preference experiment showed that treatment with MS15203 or MS15203 + Morphine produced no place preference compared to Saline indicating a lack of reward-related behavior. Taken together this data provides evidence that the novel pain therapeutic, MS15203, has minimal reward liability. Therefore, GPR171 deserves further exploration as a pain target.

**Significance Statement:** MS15203, a drug that activates the receptor GPR171, was previously shown to increase morphine analgesia. The authors use *in vivo* and histological techniques to show that it fails to activate the rodent reward circuitry, providing support for the continued exploration of MS15203 as a novel pain drug, and GPR171 a novel pain target.

## Introduction

Opioids are among the most effective pain medications available, but their addictive liability (Fields and Margolis, 2015), strong overdose potential (Pattinson, 2008; Rudd *et al*., 2016; Scholl *et al*., 2018), and limited effectiveness in the treatment of chronic pain (Volkow *et al*., 2018; Glare *et al*., 2019) necessitates the development of new pain medications (Skolnick, 2018). The opioid epidemic in the U.S.A. and its continued severity (Friedman *et al*., 2020; Silva and Kelly, 2020; Sterling and Platt, 2022), highlights the urgent need for more effective and less addictive alternatives. One area of active research addressing this need is G protein-coupled receptors (GPCR), which are the most “druggable” targets in the human body, accounting for approximately 35% of all FDA approved drugs (Wacker *et al*., 2017; Insel *et al*., 2019). However, recently deorphanized GPCRs remain an underexplored target for pain modulation. One such target is the deorphanized inhibitory GPCR, GPR171. GPR171 was discovered in 2001 (Wittenberger *et al*., 2001), and was deorphanized when its endogenous ligand was found to be the small peptide BigLEN (Gomes *et al*., 2013).

Previously our lab and others have shown GPR171 to be a promising antinociceptive target (McDermott *et al*., 2019; Cho *et al*., 2021; Ram *et al*., 2021). GPR171 is localized in the ventrolateral periaqueductal gray (McDermott *et al*., 2019) a structure essential for pain modulation and opioid action. An agonist for GPR171, MS15203, enhances morphine-mediated antinociception (McDermott *et al*., 2019), suggesting that in a clinical setting a GPR171 agonist could potentially enhance opioid analgesia, thus necessitating a lower dosage of opioid. Our lab has also shown that MS15203 is effective in reducing both inflammatory and paclitaxel-induced neuropathic pain in male, but not female, mice (Ram *et al*., 2021). Other work has shown MS15203 to effectively attenuate nociceptor mediated acute pain, inflammatory pain, and chronic constriction injury neuropathic pain (Cho *et al*., 2021). Evidence points towards GPR171 and MS15203 as a promising target and ligand for treating a wide range of pain states. However, before this receptor is further explored as a novel pain therapy, it is essential that its action on reward be assessed. It is crucial that receptor activation does not enhance morphine reward or cause reward on its own. As of yet, its role in reward, and more specifically opioid-induced reward, is unknown.

Here we explored GPR171 in rodent reward-related neural circuitry and behavior. To better understand GPR171’s role in reward we undertook the following three experiments. First, we mapped GPR171 and ProSAAS throughout the reward structures of the brain; our targeted regions of interest included the hippocampus (HPC), basolateral amygdala (BLA), nucleus accumbens (NAc), prefrontal cortex (PFC), and ventral tegmental area (VTA). Next, we measured the effect of a GPR171 agonist on VTA activation and morphine-induced activation. Lastly, we assessed the effects of GPR171 activation in an *in vivo* conditioned place preference (CPP) paradigm. Taken together, this study sought to determine GPR171’s role in reward and opioid-mediated reward for the purpose of exploring this novel receptor as a novel pain target.

## Material and Methods

### Subjects

For all three experiments, Male C57BL/6 mice (Charles Rivers Laboratories, CA), age 6-10 weeks, were used. Animals weighed 20g-35g at the start of each experiment and were given unlimited access to food and water when not undergoing experimentation. Mice were housed in groups of five in a temperature-controlled room with a 12:12 hour light:dark cycle (on at 07:00). All procedures were approved by Utah State University Institutional Care and Use Committee (Protocol #10038) and performed in accordance with the *Guide for the Care and Use of Laboratory Animals* adopted by the National Institutes of Health.

### Mapping GPR171 & ProSAAS in the reward structures of the brain

#### Immunohistochemistry

Drug naïve male C57BL/6 mice (n=5) were deeply anesthetized using isoflurane, and then transcardially perfused with 4% paraformaldehyde. Perfused brains were post-fixed for 24 hours in 4% paraformaldehyde and transferred to 1XPBS for storage and subsequent immunohistochemistry. Fixed brains were dissected for the following brain region: HPC, BLA, NAc, PFC, and VTA. Coronal sections were sliced at 50 microns with a vibrating blade microtome (Leica, VT1000S) and refrigerated in 1X PBS for storage. Immunohistochemistry was performed as described previously (McDermott *et al*., 2019) with slices washed in 1X PBS between all steps. Briefly, tissue slices were incubated for 30 minutes in 1% sodium borohydride, and subsequently placed in 5% normal goat serum blocking buffer (0.3% Triton X-100, 1X PBS) for one hour. After blocking, NAc, HPC, BLA, PFC slices were incubated and lightly shaken overnight at 4°C in 1:400 dilution of GPR171 (anti-rabbit, GeneTex, GTX108131) or 1:500 ProSAAS (anti-rabbit, MilliporeSigma ABN2268) in a solution of 0.3% Triton X-100 and 1% BSA. VTA slices were incubated in ProSAAS and GPR171 primary antibodies and a 1:500 dilution of Tyrosine Hydroxylase (TH, mouse; Invitrogen; MA1-24654). All brain region slices were then incubated for 2 hours in a 1:1000 dilution of secondary antibody: goat anti-rabbit (Alexa Flour 594, Life Technologies; A11037). VTA slices also received goat anti-mouse (Alexa Flour 488, Life Technologies; A11001). Lastly, slices were mounted and cover slipped onto glass microscope slides using ProLong Diamond Antifade (Invitrogen) mounting media. Images of fluorescent staining were captured using a Keyence BZ-X800 fluorescent microscope 4x and 20x magnification. Images were post-processed using Keyence image-analyzer software for haze reduction and lookup table contrast adjustment.

### Quantifying VTA activation after MS15203 and morphine challenge

#### Group designation and drug preparation

Animals (n=30) were randomized into four groups: Morphine (10 mg/Kg, i.p.), MS15203 (10 mg/Kg, i.p.), Morphine + MS15203 (10 mg/Kg; 10 mg/Kg, i.p.), and Saline (0.9%, i.p.). Morphine was prepared by adding stock morphine sulfate (West-Ward (Hikma) Pharmaceuticals, Eatontown, NJ) to 0.9% saline. For MS15203 (Gift from Sanjai Pathak, Queens College), solid powdered drug was dissolved in 0.9% saline. For Morphine + MS15203, powdered MS15203 was dissolved in morphine sulfate and 0.9% saline solution.

#### Experimental Design

Animals were gently restrained and interperitoneally injected with their designated drug. Ninety minutes after injection (Campos-Jurado *et al*., 2019), animals were deeply anesthetized with isoflurane, and then transcardially perfused with 4% paraformaldehyde. Perfused brains were post fixed for 24 hours in 4% paraformaldehyde and transferred to 1X PBS for storage and subsequent immunohistochemistry.

#### Immunohistochemistry

Fixed brains were dissected for VTA. Midbrain was sliced coronally at 50 microns with a vibratome and refrigerated in 1X PBS for storage. Immunohistochemistry was performed as described above. Slices were incubated overnight at 4°C in a 1:500 dilution of primary antibody: Tyrosine Hydroxylase (TH, mouse, Invitrogen; MA1-24654) and the immediate early gene protein c-Fos (c-Fos, rabbit, Abcam; ab190289). Slices were then incubated in a 1:1000 dilution of secondary antibody: goat anti-mouse (Alexa Flour 488, Life Technologies; A11001) and goat anti-rabbit (Alexa Flour 594, Life Technologies; A11037). Lastly, slices were mounted and coverslipped onto glass microscope slides using Cytoseal 60 or ProLong Diamond Antifade (Invitrogen) mounting media.

#### Microscopy and Statistical Analysis

Animals with high quality perfusion and intact VTA were selected for microscopy and statistical analysis (N=24, n= 4-8 animals per group, 2-6 slices per animal). Images of fluorescently stained VTA were obtained using a Keyence microscope at 20x magnification. 300×300 micron regions of the VTA were captured, and images were post-processed with haze reduction and contrast adjustment (lookup table settings). Red channel c-Fos activated cells were hand-counted by an experimenter blinded to experimental conditions. Statistical analysis of experimental group averages were analyzed using a one-way independent groups ANOVA and subsequent Dunnet’s multiple comparisons test with GraphPad Prism 9 software.

### Assessment of reward of GPR171 ligands *in vivo* using conditioned place preference

#### Drug designations

Animals (n=48, 10-14 animals per group) were randomly divided into four groups as above: Morphine (10mg/Kg), Morphine + MS15203 (10mg/Kg + 10mg/Kg), MS15203 (10mg/Kg), or Saline. Animals were gently restrained and administered their designated treatment interperitoneally with a 27-gauge hypodermic needle at a volume of 10mL/Kg. All drugs were prepared as described above.

#### Conditioned place preference experimental design

The rodent behavioral assay, conditioned place preference (CPP) was used to assess the *in vivo* propensity of MS15203 to invoke reward or aversion alone, and with morphine. The CPP apparatus (Maze Engineers) consisted of two distinct chambers (34 × 25 cm) separated by a neutral middle compartment (6 × 25 cm). The two chambers, one grey-walled and one striped-walled, were distinct in smell and floor texture (see Figure 6). Between trials the grey chamber was cleaned with a mixture of water and dish soap while the striped chamber was cleaned with ethanol. The neutral middle compartment was white-walled and free of smell. The experimental paradigm lasted 10 days, with conditioning days on Days 2-9, and a pretest on Day 1 and posttest on Day 10. For conditioning days, mice were given their designated drug or saline and placed in the grey or striped chamber, with the inability to access other compartments, for 30 minutes. On odd days (e.g., 3, 5, 7, 9) animals received their designated drug while on alternate even days (e.g., 2, 4, 6, 8) all animals received saline. The chamber that animals received saline or drug were randomly counterbalanced to account for innate preference for an experimental chamber. The control group, Saline, received saline in both chambers on alternate days. On Day 10 (posttest) mice were placed in the middle compartment and allowed 15 minutes to freely move between all compartments. Time spent in each compartment was quantified.

#### Data collection and analysis

Time spent in each chamber of the CPP apparatus was visualized and measured using an ANY-maze video camera and motion tracking software. Group averages were compared using an omnibus one-way independent groups ANOVA followed by a Dunnet’s *post hoc* multiple comparisons. All analysis was done using GraphPad Prism 9 software.

## Results

### GPR171 and ProSAAS are localized throughout reward-related structures

Immunohistochemistry results show expression of GPR171 throughout four reward-related structures of the mouse brain: HPC, BLA, NAc, PFC (Figure 1). Similarly, ProSAAS is found in all four structures: HPC, BLA, NAc, and PFC (Figure 2), but there was markedly less localization of ProSAAS in the BLA than surrounding tissue. GPR171 appeared to be localized in dopamine neurons of the VTA as observed by yellow-orange staining in the GPR171-TH overlay (Figure 3b). Notably GPR171 punctae appeared to be localized largely in the cell bodies of dopamine neurons (Figure 3c). In contrast, ProSAAS appeared to be primarily localized outside of dopamine neurons of the VTA (Figure 4b) and appeared to show little colocalization between the two channels. In total, these results suggest the presence, but differential expression of, GPR171 and ProSAAS within the VTA.

**Figure 1.**
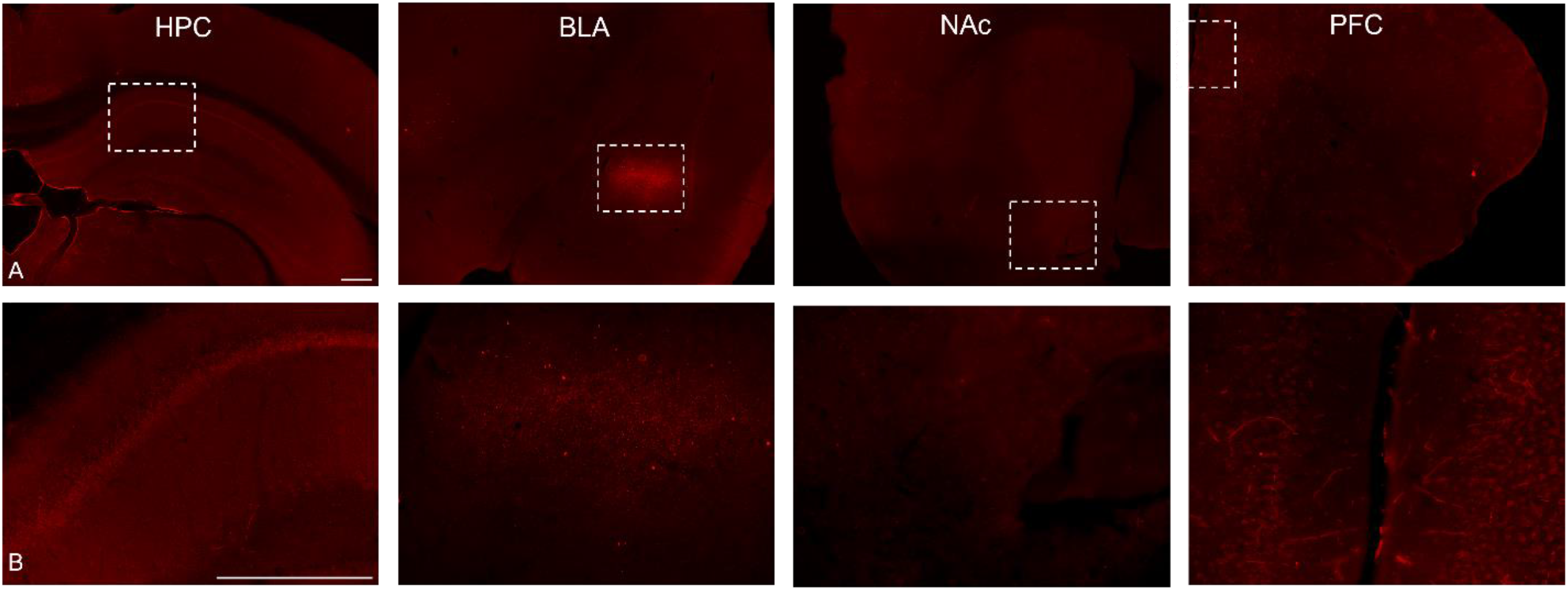
Localization of GPR171 in reward and addiction-related structures of the brain. 4x (A) and 20x (B) images of the hippocampus (HPC), basolateral amygdala (BLA), nucleus accumbens (NAc), and prefrontal cortex (PFC). All four regions show GPR171 expression. In particular, GPR171 is distinctly localized in the BLA. Scale bars = 300μm

**Figure 2.**
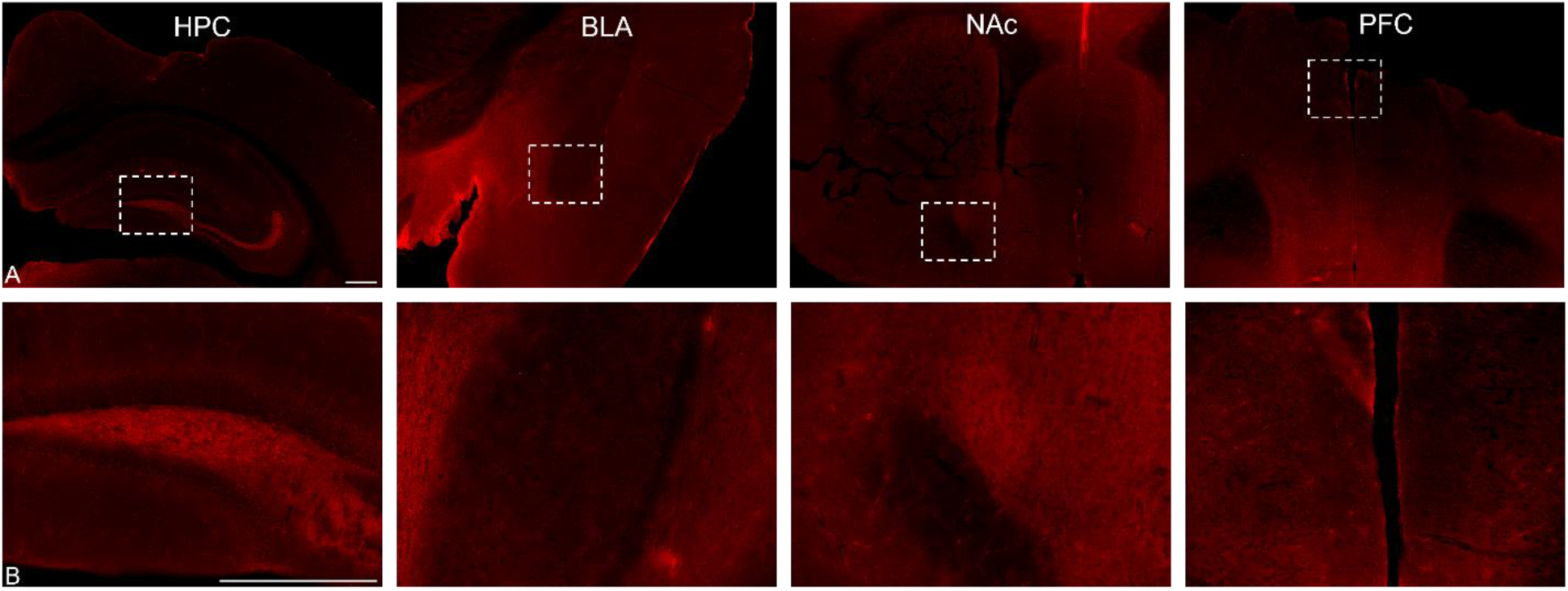
Localization of ProSAAS in reward and addiction-related structures of the brain. 4x (A) and 20x (B) images of the hippocampus (HPC), nucleus accumbens (NAc), and prefrontal cortex (PFC). All regions show ProSAAS localization. In particular, ProSAAS shows less expression in the BLA than its surrounding tissue. Scale bars = 300μm

**Figure 3.**
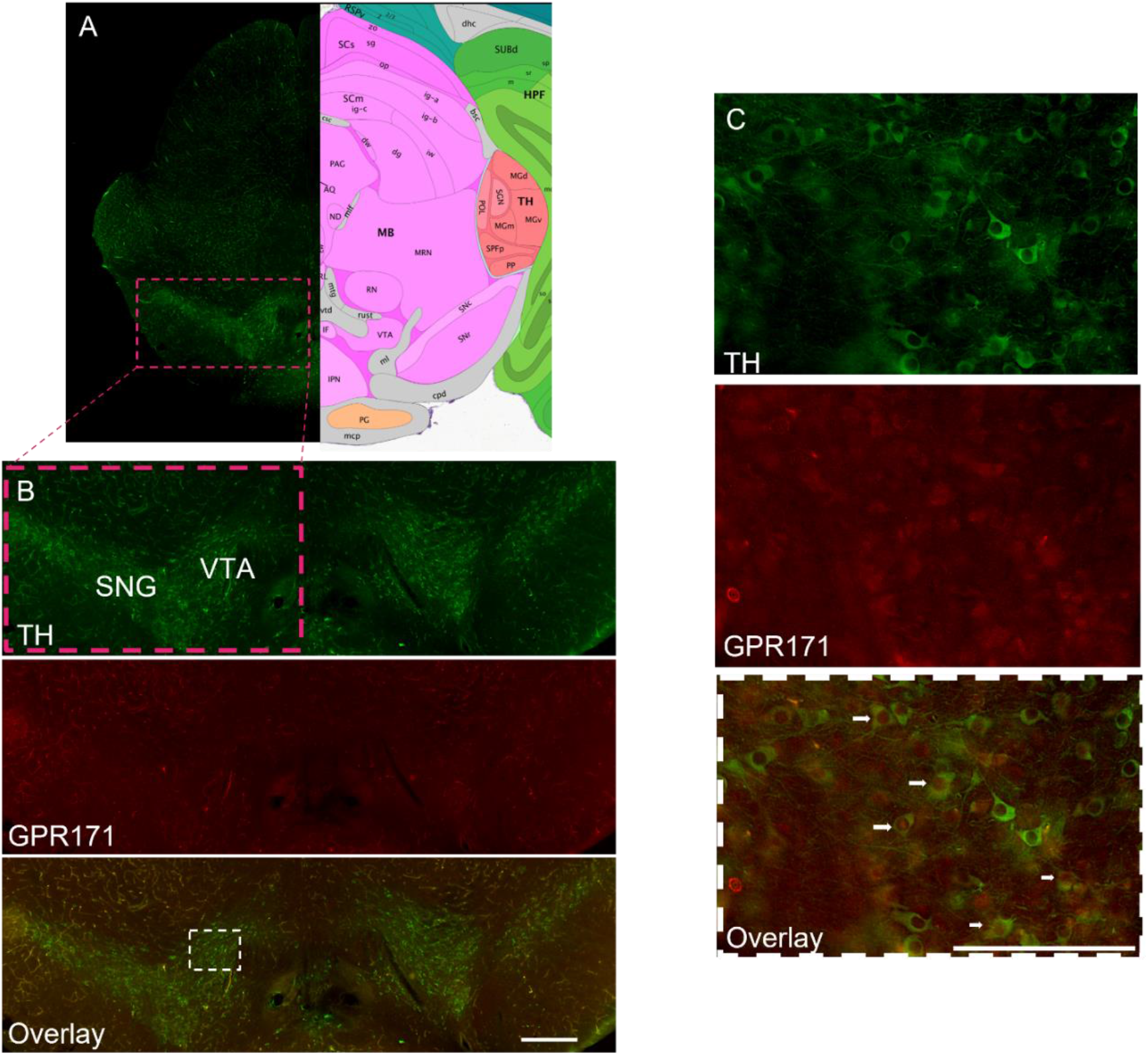
Expression of GPR171 in dopamine neurons of the ventral tegmental area (A) A midbrain coronal section showing dopamine neurons of the left ventral tegmental area (VTA) and substantia nigra (SNG), and a schematic adapted from the Allen Brain Atlas’ Mouse Brain Reference Atlas (Mouse P56 Coronal) showing, among other regions, right VTA and SNG. (B) GPR171 expression in the VTA and SNG left and right VTA and SNG stained for TH, GPR171, and an overlay of the two channels show GPR171 colocalized in subset of dopamine neurons. (C) a 20x capture of left VTA from white box in 3B showing GPR171 staining. Arrows on the TH-GPR171 overlay show regions of colocalization, mainly in the cell body of TH neurons. Scale bars = 300μm

**Figure 4.**
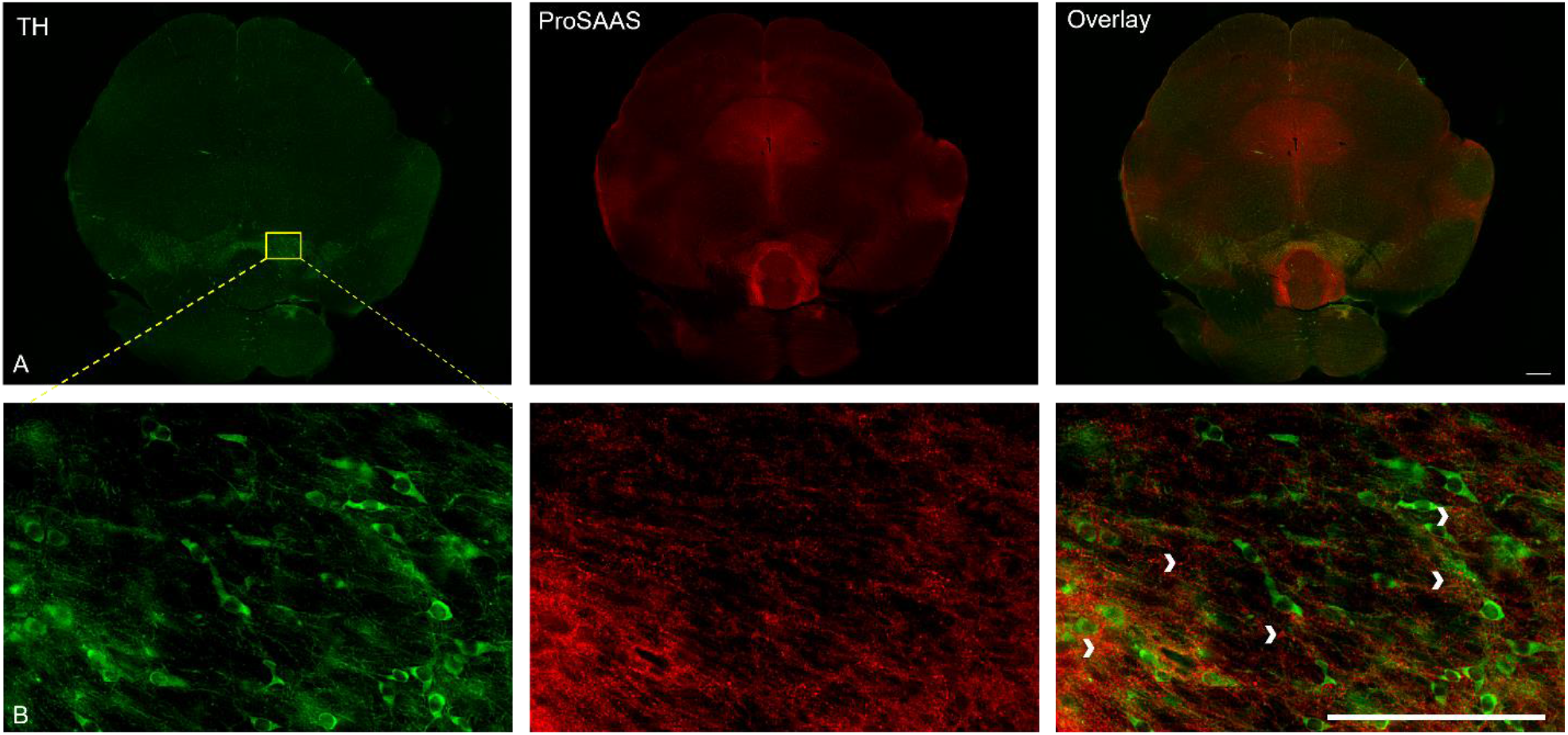
ProSAAS localization in the VTA (A) A stitched 4x coronal midbrain section displaying TH and ProSAAS staining (B) 20x image of right VTA showing TH and ProSAAS staining. Chevrons display ProSAAS punctae located outside of dopamine neurons. Scale bars = 300μm

### MS15203 does not increases c-Fos expression in the VTA

IHC was performed on midbrain sections stained for TH and c-Fos for four experimental groups: Saline, MS15203, Morphine, and Morphine + MS15203 (Figure 5). c-Fos activated cells in the VTA were quantified and group differences were analyzed using a one-way independent groups ANOVA. An overall main effect was statistically significant [F(3,55)=11.37, *p*<.0001]. Since there was an overall main effect, a Dunnet’s *post hoc* multiple comparisons was used to compare group means to Saline. Unsurprisingly, Morphine and Saline showed a significant difference (Dunnet’s, p=0.0165). Morphine + MS15203 and Saline were significantly different from each other (Dunnet’s, *p*<0.0001). Importantly, there was no significant difference between MS15203 and Saline (Dunnet’s, *p*=0.9971).

**Figure 5.**
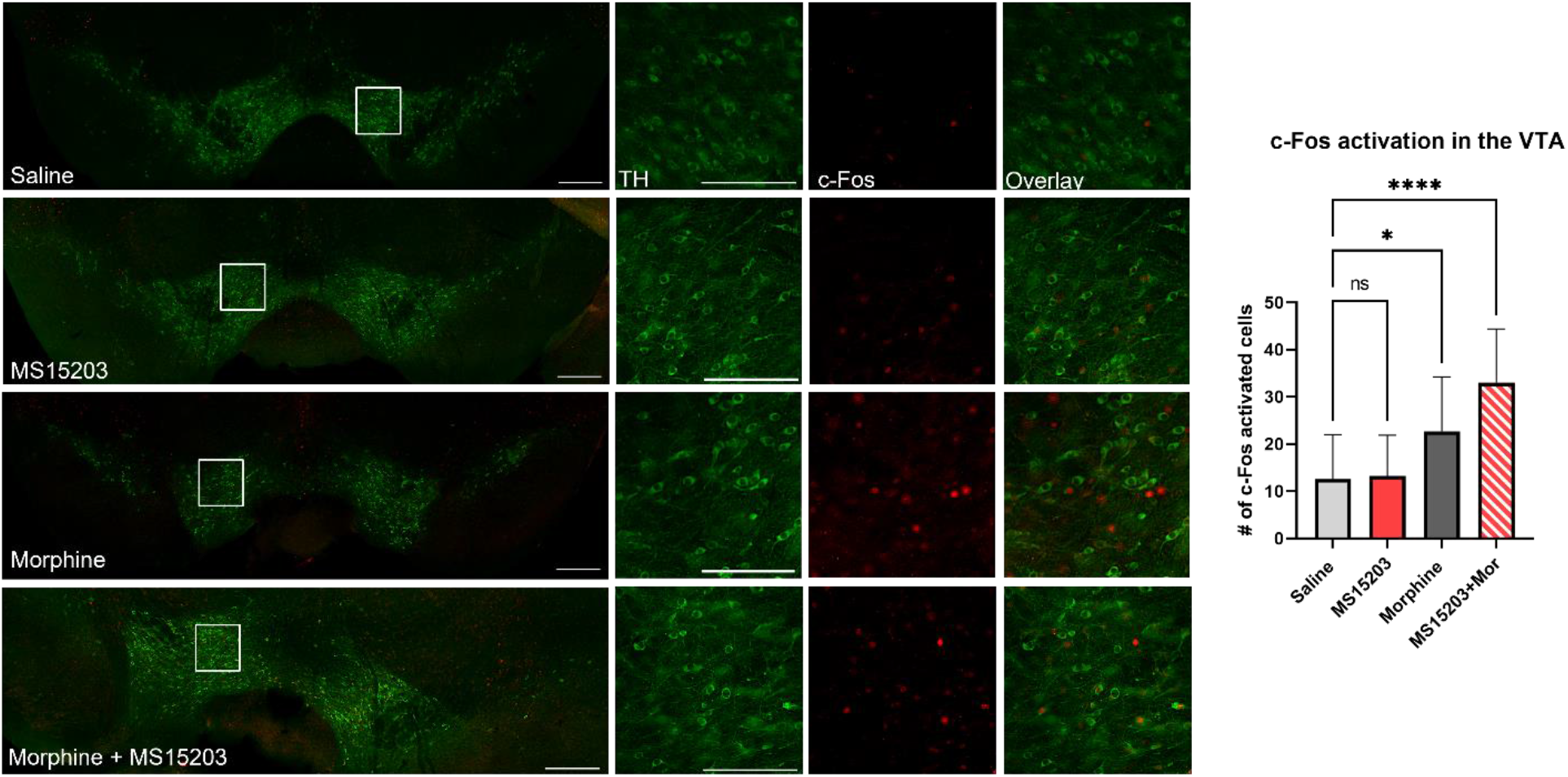
Quantification of c-Fos activated cells in the VTA following drug treatment. Saline, MS15203, Morphine, and Morphine + MS15203 show the overlay of c-Fos and TH captured at 20x and stitched 8×3. White boxes delineate the 300×300 region of VTA used for c-Fos quantification and which are respectively showcased to the right of each stitched image. 300×300 images show TH, c-Fos and overlay for their respective group. Ninety minutes after designated drug treatment (Saline; 0.9%, MS15203; 10 mg/Kg, Morphine; 10 mg/Kg, Morphine + MS15203; 10 mg/Kg, 10 mg/Kg) animals were transcardially perfused with paraformaldehyde. Number of c-Fos activated cells in the VTA were quantified. A one-way independent groups ANOVA with Dunnet’s multiple comparisons was run to analyze group averages. MS15203 (n=5, slices =14) was compared to Saline (n=8, slices =18) (p=0.997). Morphine (n=7, slices =16) and Morphine + MS15203 (n=4, slices=11) were also compared to Saline (p = 0.0165, p <.001, respectively). Scale bars = 300μm

### MS15203 fails to induce place preference or increase morphine place preference

The behavioral conditioned place preference (CPP) paradigm included the groups: Saline, Morphine, MS15203, and MS15203 + Morphine (Figure 6). A one-way independent groups ANOVA on group averages showed an overall omnibus effect [*F*(3,44)= 5.252, *p*=0.0035]. Since there was an overall main effect, a Dunnet’s *post hoc* multiple comparisons was used to compare group means to Saline. Unsurprisingly, Morphine vs. Saline were statistically significant from one another (Dunnet’s, *p*=0.0280). Remarkably, there was no significant difference between MS15203 + Morphine and Saline (Dunnet’s, *p*=0.0717), and importantly, no statistical difference between MS15203 and Saline (Dunnet’s, *p*=0.9251).

**Figure 6.**
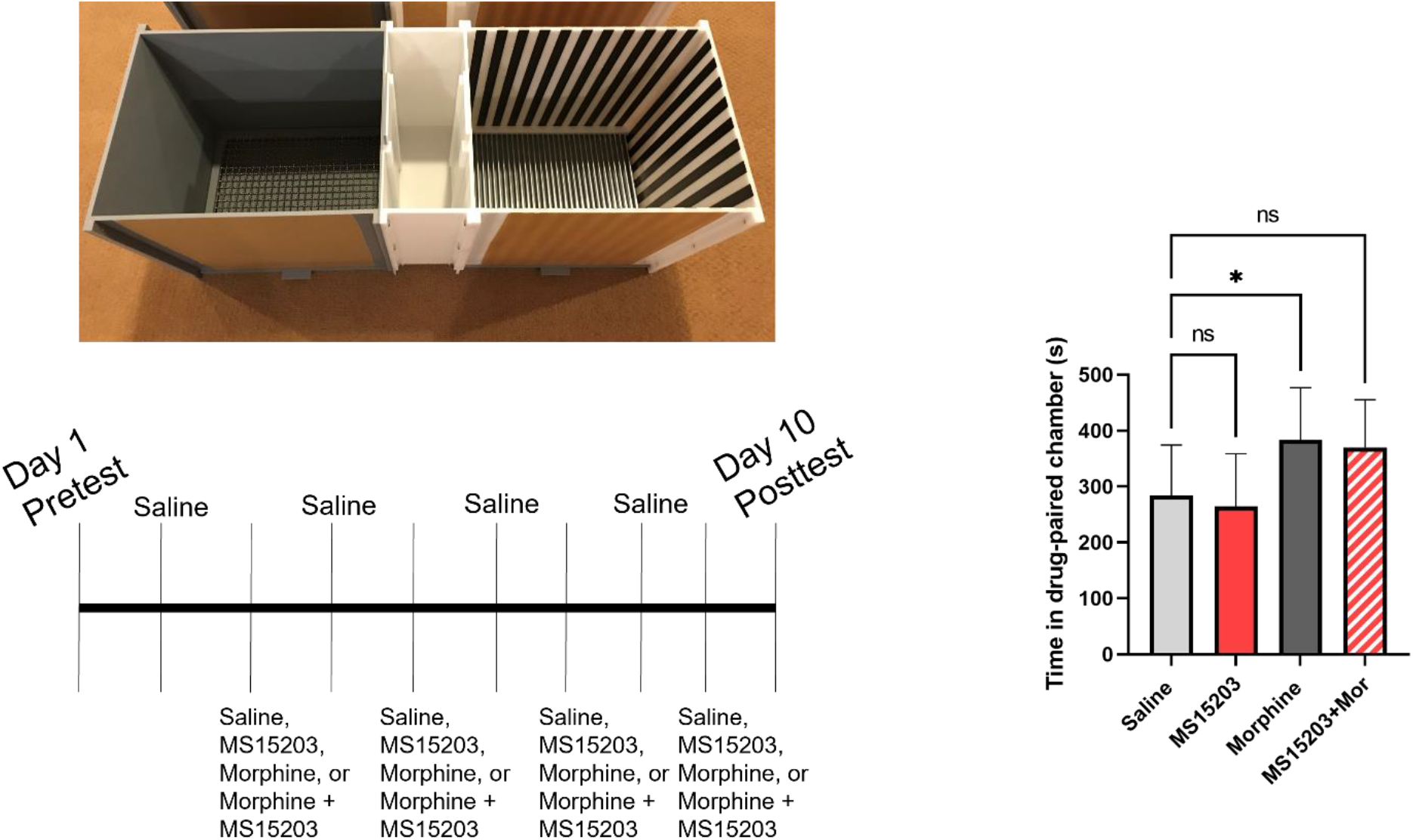
Assessment of Conditioned place preference. Time spent in drug paired chamber on Day 10 for each experimental group were compared using a one-way independent groups ANOVA. Saline (n =10) and MS1520 (n=11) show the lowest preference scores and were compared using a Dunnet’s multiple comparison (p =0.9251). Morphine +MS15203 (n=13) and Morphine (n=14) were compared to Saline (p=0.0717, p=0.0280, respectively).

## Discussion

In this study we showed that ProSAAS and GPR171 are found in important reward related brain structures: hippocampus, basolateral amygdala, nucleus accumbens, prefrontal cortex, and ventral tegmental area. While GPR171 is localized in a subset of dopamine neurons within the VTA, minimal localization of ProSAAS is found within dopamine neurons of the VTA. Also using fluorescent IHC we observed that MS15203 does not alter c-Fos activation in the VTA. Similarly, behavioral data in our conditioned place preference experiment showed that MS15203 administration failed to induce place preference alone and failed to increase morphine-induced place preference. Taken together, these studies lend credence for MS15203 as a novel pain therapeutic and enhancer of morphine analgesia without reward.

Previously, our lab showed that MS15203 increases morphine antinociception likely mediated through the actions of the periaqueductal grey (McDermott *et al*., 2019). The current study is the first to evaluate the role of GPR171 in drug reward. Peripherally related work has been on the topic of ProSAAS and reward. For example, it has been shown that in response to chronic cocaine administration, ProSAAS becomes down-regulated in the VTA and the NAc, though ProSAAS knockouts still show place preference to cocaine (Berezniuk *et al*., 2017). Another peptide derived from ProSAAS, PEN, the endogenous ligand for the deorphanized receptor, GPR83, has been explored in reward (Fakira *et al*., 2019). GPR83 is expressed in the VTA and GPR83 knockdown in the NAc results in decreased morphine place preference in male rodents. The NAc is also important motivation for non-drug stimuli. BigLEN reduces NAc transmission, and antagonizing the GPR171 receptor results in decreased persistent food seeking (Smith *et al*., 2022). These studies leave open the question as to whether GPR171 activation mediates reward and VTA engagement.

Behaviorally, the current study showed that MS15203 and MS15203 + Morphine failed to induce reward measured by the conditioned place preference assay. Previous studies have linked alterations in morphine CPP to actions through dopamine receptors. One study has shown that a D1 receptor antagonist blocks morphine place preference in rats (Grenier *et al*., 2022). Additionally, the atypical antipsychotic quetiapine, which also has antagonistic properties at the dopamine receptors, and at the serotonin receptors, attenuates morphine CPP in rats (Khezri *et al*., 2022). Similar to GPR171, Neuropeptide S, the endogenous ligand for the previously orphaned GPCR, GPR154, has been shown to be neutral alone, but decrease morphine place preference in mice (Li *et al*., 2009). Additionally, it has been shown to cause antinociception when administered nasally (Medina *et al*., 2014) and intracerebroventricularly (Peng *et al*., 2010).

In large part, this study focused on the relationship between GPR171 and the ventral tegmental area. The VTA is a major dopaminergic center of the brain, and release of dopamine from this area into the NAc is a crucial mechanism for opioid-induced reward (Fields and Margolis, 2015; Kim *et al*., 2016) and reward generally. Opioid agonists disinhibit GABA interneurons that tonically suppress dopamine release in the VTA resulting in activation of dopamine neurons (Johnson and North, 1992; Trang *et al*., 2015). Though the mechanism of VTA activation varies depending on the drug involved, reward is largely mediated by dopamine release in structures like the NAc, regardless of substance (Fujita *et al*., 2019). In the current study IHC staining for c-Fos was used as an indirect indicator of neuronal activation in the VTA as has been done previously (Dela Cruz *et al*., 2016; Dehkordi *et al*., 2017; Campos-Jurado *et al*., 2019). Indeed, the rapid transcription from neuronal activation has made c-Fos one of the most frequently utilized immediate early genes for locating neuronal activation in addiction research (Cruz *et al*., 2015).

While the Morphine + MS15203 treated mice were not significant from saline treated in the CPP experiment, this group was significant from saline in the c-Fos experiment. In other words, though Morphine + MS15203 does not result in reward-related behavior, it does result in a significant VTA activation compared to saline. One possible explanation for why MS15203 increases activation of a major reward structure, but fails to induce behavioral reward, may be the localization of GPR171. First of all, GPR171 is localized on dopamine neurons and as an inhibitory GPCR would result in diminished dopaminergic activity. Secondly, GPR171 may be primarily localized on a subset of GABA neurons in the VTA (Fields and Margolis, 2015; Yu *et al*., 2019; Galaj *et al*., 2020) that synapse onto non-dopamine postsynaptic neurons. GPR171 agonist would produce disinhibition of these GABA neurons that specifically synapse onto non-dopamine neurons of the VTA and would appear as increased activation, but without producing reward related behaviors associated with dopamine actions. The exact mechanism is unclear, but since the VTA is a heterogenous region (Fields and Margolis, 2015) and contains both glutamate and GABA neurons the mechanism likely involves multiple cell types. Indeed, we have shown that GPR171 is expressed in GABA and glutamate neurons of the PAG and BLA (Bobeck *et al*., 2017; McDermott *et al*., 2019), but it is unknown what other cell types besides dopamine express GPR171 within the VTA. To determine the exact relationship between dopamine and GPR171 future studies should look at the percentage of VTA dopamine neurons that show MS15203-activation compared to non-dopamine neurons of the VTA.

The presence of both ProSAAS and GPR171 in the ventral tegmental area suggests an endogenous role for the BigLEN-GPR171 neuropeptide system in this brain structure. Interestingly, IHC and fluorescent microscopy demonstrated that GPR171 and ProSAAS are differentially dispersed throughout the VTA although GPR171 appeared to be located primarily in the cell bodies of dopamine neurons, and ProSAAS was distinctly located outside of these structures. ProSAAS-containing neurons are likely secreting BigLEN presynaptically onto the dopamine neurons containing GPR171. In this scenario, the BigLEN-GPR171 system would likely be acting endogenously to inhibit dopamine release, which is congruous with our observation that MS15203 shows minimal place preference. It is also possible that the high ProSAAS expression points towards a larger role being played by the other small ProSAAS-derived peptides, like PEN, the endogenous ligand for GPR83. However, since ProSAAS is one of the most highly expressed proteins in the brain (Fricker, 2010) It’s possible that ProSAAS is simply acting in this region similar to its theorized house-keeping role as a chaperone protein (Hoshino *et al*., 2014).

Intriguingly, we also showed differential expression of ProSAAS and GPR171 in the basolateral amygdala. Our lab has shown similar high expression of GPR171 in this region in the past and knockdown of GPR171 in this region lowers anxiety (Bobeck *et al*., 2017), but to date no one has looked at ProSAAS expression in the BLA. While GPR171 showed high expression in the BLA, ProSAAS showed markedly low expression compared to surrounding amygdala regions, such as the central amygdala. Future studies will look at the behavioral effect of knocking down ProSAAS in this region and the naïve colocalization between ProSAAS and GPR171 to better understand the actions of BigLEN-GPR171 system in the amygdala.

In conclusion, we have shown that MS15203 does not increase c-Fos VTA activation and it fails to induce place preference in mice *in vivo*. The current data lends evidence and confidence towards the continued exploration of GPR171 as a novel pain therapeutic, capable of enhancing opioid analgesia without increasing its reward. Future studies should probe the exact mechanism regulating GPR171 actions in the VTA and further explore the relationship between ProSAAS and GPR171 in the mesolimbic pathway. Further assessment of the safety profile of MS15203 should be investigated including side effects such as withdrawal, tolerance and opioid-induced respiratory depression.

## List of abbreviations

BLA: Basolateral Amygdala
CPP: Conditioned Place Preference
HPC: Hippocampus
NAc: Nucleus Accumbens
PFC: Prefrontal Cortex
TH: Tyrosine Hydroxylase
VTA: Ventral Tegmental Area

## Acknowledgements

We would like to thank Dr. Sanjai Pathak from Queens college, NY for his generous gift of MS15203.

## Footnotes

This work was funded on startup funds from Utah State University (to ENB) and a Pharmacology & Toxicology Startup grant from the PhRMA Foundation (to ENB) and NIH grant (TR003667-01).

## Contribution

Participating in research design: McDermott, Ram, Bobeck,

Conducted experiments: McDermott, Ram, Mattoon, Haderlie, Raddatz, Thomason, Bobeck

Wrote or contributed to the writing of the manuscript: McDermott, Ram, Mattoon, Haderlie, Raddatz, Bobeck

## References

Berezniuk I, Rodriguiz RM, Zee ML, Marcus DJ, Pintar J, Morgan DJ, Wetsel WC, and Fricker LD (2017) ProSAAS-derived peptides are regulated by cocaine and are required for sensitization to the locomotor effects of cocaine. J Neurochem 143:268, NIH Public Access.

Bobeck EN, Gomes I, Pena D, Cummings KA, Clem RL, Mezei M, and Devi LA (2017) The BigLEN-GPR171 Peptide Receptor System Within the Basolateral Amygdala Regulates Anxiety-Like Behavior and Contextual Fear Conditioning. Neuropsychopharmacol 2017 4213 42:2527–2536, Nature Publishing Group.

Campos-Jurado Y, Igual-Lopez M, Padilla F, Zornoza T, Granero L, Polache A, Agustin-Pavon C, and Hipolito L (2019) Activation of MORs in the VTA induces changes on cFos expression in different projecting regions: Effect of inflammatory pain. Neurochem Int 131:104521, England.

Cho PS, Lee HK, Choi YI, Choi SI, Lim JY, Kim M, Kim H, Jung SJ, and Hwang SW (2021) GPR171 Activation Modulates Nociceptor Functions, Alleviating Pathologic Pain. Biomedicines 9:256, DPI.

Cruz FC, Javier Rubio F, and Hope BT (2015) Using c-fos to study neuronal ensembles in corticostriatal circuitry of addiction. Brain Res 1628:157–173.

Dehkordi O, Rose JE, Dávila-García MI, Millis RM, Mirzaei SA, Manaye KF, and Jayam-Trouth A (2017) Neuroanatomical Relationships between Orexin/Hypocretin-Containing Neurons/Nerve Fibers and Nicotine-Induced c-Fos-Activated Cells of the Reward-Addiction Neurocircuitry. J Alcohol drug Depend 5:273.

Dela Cruz JAD, Coke T, and Bodnar RJ (2016) Simultaneous Detection of c-Fos Activation from Mesolimbic and Mesocortical Dopamine Reward Sites Following Naive Sugar and Fat Ingestion in Rats. J Vis Exp 53897, MyJove Corporation.

Fakira AK, Peck EG, Liu Y, Lueptow LM, Trimbake NA, Han M-H, Calipari ES, and Devi LA (2019) The role of the neuropeptide PEN receptor, GPR83, in the reward pathway: Relationship to sex-differences. Neuropharmacology 157:107666.

Fields HL, and Margolis EB (2015) Understanding opioid reward. Trends Neurosci 38:217–225, Elsevier Ltd.

Fricker LD (2010) Analysis of mouse brain peptides using mass spectrometry-based peptidomics: Implications for novel functions ranging from non-classical neuropeptides to microproteins.

Friedman SR, Krawczyk N, Perlman DC, Mateu-Gelabert P, Ompad DC, Hamilton L, Nikolopoulos G, Guarino H, and Cerdá M (2020) The Opioid/Overdose Crisis as a Dialectics of Pain, Despair, and One-Sided Struggle. Front public Heal 8:540423, Frontiers Media S.A.

Fujita M, Ide S, and Ikeda K (2019) Opioid and nondopamine reward circuitry and state-dependent mechanisms. Ann N Y Acad Sci 1451:29–41, United States.

Galaj E, Han X, Shen H, Jordan CJ, He Y, Humburg B, Bi G-H, and Xi Z-X (2020) Dissecting the Role of GABA Neurons in the VTA versus SNr in Opioid Reward. J Neurosci 40:8853–8869.

Glare P, Aubrey KR, and Myles PS (2019) Transition from acute to chronic pain after surgery. Lancet 393:1537–1546.

Gomes I, Aryal DK, Wardman JH, Gupta A, Gagnidze K, Rodriguiz RM, Kumar S, Wetsel WC, Pintar JE, Fricker LD, and Devi LA (2013) GPR171 is a hypothalamic G protein-coupled receptor for BigLEN, a neuropeptide involved in feeding. Proc Natl Acad Sci 110:16211–16216.

Grenier P, Mailhiot MC, Cahill CM, and Olmstead MC (2022) Blockade of dopamine D1 receptors in male rats disrupts morphine reward in pain naïve but not in chronic pain states. J Neurosci Res 100:297–308, United States.

Hoshino A, Helwig M, Rezaei S, Berridge C, Eriksen JL, and Lindberg I (2014) A novel function for proSAAS as an amyloid anti-aggregant in Alzheimer’s disease. J Neurochem 128:419–430, England.

Insel PA, Sriram K, Gorr MW, Wiley SZ, Michkov A, Salmerón C, and Chinn AM (2019) GPCRomics: An Approach to Discover GPCR Drug Targets. Trends Pharmacol Sci 40:378–387.

Johnson SW, and North RA (1992) Opioids excite dopamine neurons by hyperpolarization of local interneurons. J Neurosci 12:483–488.

Khezri A, Mohsenzadeh MS, Mirzayan E, Bagherpasand N, Fathi M, Abnous K, Imenshahidi M, Mehri S, and Hosseinzadeh H (2022) Quetiapine attenuates the acquisition of morphine-induced conditioned place preference and reduces ERK phosphorylation in the hippocampus and cerebral cortex. Am J Drug Alcohol Abuse 1–11, England.

Kim J, Ham S, Hong H, Moon C, and Im H-I (2016) Brain Reward Circuits in Morphine Addiction. Mol Cells 39:645–53.

Li W, Gao Y-H, Chang M, Peng Y-L, Yao J, Han R-W, and Wang R (2009) Neuropeptide S inhibits the acquisition and the expression of conditioned place preference to morphine in mice. Peptides 30:234–240.

McDermott M V, Afrose L, Gomes I, Devi LA, and Bobeck EN (2019) Opioid-induced signaling and antinociception are modulated by the recently deorphanized receptor, GPR171. J Pharmacol Exp Ther 371:56–62, American Society for Pharmacology and Experimental Therapeutics.

Medina G, Ji G, Grégoire S, and Neugebauer V (2014) Nasal application of neuropeptide S inhibits arthritis pain-related behaviors through an action in the amygdala. Mol Pain 10:32.

Pattinson KTS (2008) Opioids and the control of respiration. Br J Anaesth 100:747–758, England.

Peng Y-L, Zhang J-N, Chang M, Li W, Han R-W, and Wang R (2010) Effects of central neuropeptide S in the mouse formalin test. Peptides 31:1878–1883.

Ram A, Edwards T, McCarty A, Afrose L, McDermott M V., and Bobeck EN (2021) GPR171 Agonist Reduces Chronic Neuropathic and Inflammatory Pain in Male, but not in Female Mice. bioRxiv 2021.04.16.440030, Cold Spring Harbor Laboratory.

Rudd RA, Seth P, David F, and Scholl L (2016) Increases in Drug and Opioid-Involved Overdose Deaths - United States, 2010-2015. MMWR Morb Mortal Wkly Rep 65:1445–1452, United States.

Scholl L, Seth P, Kariisa M, Wilson N, and Baldwin G (2018) Drug and Opioid-Involved Overdose Deaths - United States, 2013-2017. MMWR Morb Mortal Wkly Rep 67:1419–1427.

Silva MJ, and Kelly Z (2020) The escalation of the opioid epidemic due to COVID-19 and resulting lessons about treatment alternatives. Am J Manag Care 26:e202–e204, United States.

Skolnick P (2018) The Opioid Epidemic: Crisis and Solutions. Annu Rev Pharmacol Toxicol 58:143–159, United States.

Smith NK, Plotkin JM, and Grueter BA (2022) Hunger dampens a nucleus accumbens circuit to drive persistent food seeking. Curr Biol 32:1689–1702.e4, Elsevier.

Sterling P, and Platt ML (2022) Why Deaths of Despair Are Increasing in the US and Not Other Industrial Nations—Insights From Neuroscience and Anthropology. JAMA Psychiatry 79:368–374.

Trang T, Al-Hasani R, Salvemini D, Salter MW, Gutstein H, and Cahill CM (2015) Pain and Poppies: The Good, the Bad, and the Ugly of Opioid Analgesics. J Neurosci 35:13879–13888, Society for Neuroscience.

Volkow N, Benveniste H, and McLellan AT (2018) Use and Misuse of Opioids in Chronic Pain. Annu Rev Med 69:451–465, United States.

Wacker D, Stevens RC, and Roth BL (2017) How Ligands Illuminate GPCR Molecular Pharmacology. Cell 170:414–427.

Wittenberger T, Schaller HC, and Hellebrand S (2001) An expressed sequence tag (EST) data mining strategy succeeding in the discovery of new G-protein coupled receptors. J Mol Biol 307:799–813, England.

Yu X, Li W, Ma Y, Tossell K, Harris JJ, Harding EC, Ba W, Miracca G, Wang D, Li L, Guo J, Chen M, Li Y, Yustos R, Vyssotski AL, Burdakov D, Yang Q, Dong H, Franks NP, and Wisden W (2019) GABA and glutamate neurons in the VTA regulate sleep and wakefulness. Nat Neurosci 22:106–119.

